# Multimodal characterization and optogenetic potential of the bistable G_i/o_-coupled vertebrate ancient opsin from the flashlight fish *Anomalops katoptron*

**DOI:** 10.1101/2025.02.20.639284

**Authors:** Lennard Rohr, Daniel Zhornist, Ori Bermann, Christiane Ruse, Marius Seidenthal, Genevieve Marie Auger, Caroline Naber, Alina Kirchhof, Caroline I. Güers, Melanie D. Mark, Till Rudack, Peter Soba, Alexander Gottschalk, Moran Shalev-Benami, Ida Siveke, Stefan Herlitze

## Abstract

Vertebrate ancient long opsin, or VAL opsin, is a light-sensitive protein that is found within and outside the visual system in vertebrates. In accordance with its wide distribution in the retina, brain, testis and skin, VAL is suggested to play a role in light-dependent physiological processes that are beyond vision. However, many aspects of the physiological properties and specific functions of VAL remain unclear. Here we identified and characterized the VAL opsin from the flashlight fish *Anomalops katoptron* (*_Ak_*VAL) and show that this opsin is bistable and reversibly converts between active and inactive states by responding to cycles of green and blue/UV lights. We further show that *_Ak_*VAL couples to the G_i/o_ pathway and controls the activity of GIRK channels in a bistable manner. In line with this, we demonstrated that *_Ak_*VAL modulates neuronal activity in cerebellar Purkinje cells, where neuronal activity is reduced by UV/blue light and increased by green/red light illumination. In addition, upon the *in vivo* expression of *_Ak_*VAL in neurons innervating body muscles of *Caenorhabditis elegans* the worm’s body movement can be bidirectionally controlled altering blue/UV and green illuminations. These data highlight the potential of *_Ak_*VAL as an optogenetic tool to control cells *in vitro* and *in vivo*, in a bistable manner.

## Introduction

The ability to control biological processes with light has revolutionized our understanding of complex neuronal systems (Emiliani et al. 2022; Boyden et al. 2005; Li et al. 2005; Nagel et al. 2005). The rapidly evolving field of optogenetics uses the power of naturally or genetically engineered light-sensitive proteins to enable precise spatiotemporal control of cellular signaling to understand neuronal network function (Rost et al. 2022; Karapinar et al. 2021; Rajasethupathy et al. 2016). The disadvantages of traditional pharmaceutical approaches for the treatment of neurological disorders, such as limited spatiotemporal resolution and off-target effects, might be overcome by new optogenetic-based therapies (Urban und Roth 2015; Mühlhäuser et al. 2017; Stuber und Mason 2013; Xie et al. 2013; Zhang und Cohen 2017; Abreu und Levitz 2020).

Among the diverse range of light-sensitive proteins, G protein-coupled receptor (GPCR)-based opsins have emerged as promising candidates for optogenetic applications (Abreu und Levitz 2020). Compared to ionotropic inhibitory tools, these opsins can modulate a wider range of downstream signaling cascades (Liu et al. 2024). Moreover, GPCR-based tools are characterized by their sensitivity to low intensity light (Berry et al. 2019; Karapinar et al. 2021; Wagdi et al. 2022; Masseck et al. 2014) and their possibility to be switched on and off by two different wavelengths of light in case of bistable or multistable opsins (Eickelbeck et al. 2020; Mahn et al. 2021; Spoida et al. 2016; Surdin et al. 2023; Wietek et al. 2024).

Optogenetic neuronal excitation has been extensively studied and successfully applied with a variety of tools (Li et al. 2005; Gauvain et al. 2021). However, the development of inhibitory tools has been a challenging process, with significant advancements only emerging in recent years (Wiegert et al. 2017). Common early tools such as halorhodopsin (Han und Boyden 2007) and archaerhodopsin (Chow et al. 2010) have been widely used to silence neuronal activity by chloride or proton-induced hyperpolarization, but require high light intensities and depend on precise ion concentrations that can vary in between cell types or developmental states (Kaila et al. 2014; Pugh und Jahr 2011). Although tools such as chloride-gated GtACRs (Govorunova et al. 2015; Li et al. 2019) or the newly discovered light-dependent potassium channels (Vierock et al. 2022; Govorunova et al. 2022) with improved kinetics, increased light sensitivity or distinct ion specificity, overcome many of these limitations. However, GPCR-based inhibitory tools offer further unique prospects of neuronal silencing by activating native signaling cascades as has been first shown for vertebrate rhodopsin (Li et al. 2005).

While tools such as Rh1 (Li et al. 2005; Oh et al. 2010), Opn5 (Wagdi et al. 2022), parapinopsin (Eickelbeck et al. 2020; Copits et al. 2021), SWO/LWO (Masseck et al. 2014), mosquito Opn3 (Mahn et al. 2021), and PdCo (Wietek et al. 2024) have been successfully used to inhibit neuronal activity, the wide range of diverse application requires improved tools with respect to light sensitivity, spectral properties, kinetics, targeting or G protein pathway specificity. The creation or discovery of bistable opsins is of particular interest because of the feasibility of light-dependent activation and deactivation (Tsukamoto und Terakita 2010) and the induction of long-lasting effects after only brief light pulses, bypassing potential phototoxic effects.

In this study, we characterize the **v**ertebrate **a**ncient **l**ong opsin (VAL) found in the retina of the flashlight fish ***A****nomalops **k**atoptron* (*_Ak_*VAL). VAL opsins are GPCR photoreceptors that represent a distinct class of light-sensitive proteins and were originally discovered in the retina of various fish species (Soni und Foster 1997; Soni et al. 1998; Jenkins et al. 2003; Kojima et al. 2000). Later, these opsins were also found in the brain and spinal cord of fish (Kojima et al. 2008; Philp et al. 2000; Davies et al. 2012), amphibians (Sato et al. 2011), and various regions of the avian brain (Tomonari et al. 2007; Halford et al. 2009). VAL opsins have been implicated in photoperiodic behavior (Halford et al. 2009; Davies et al. 2012; García-Fernández et al. 2015), neuroendocrine regulation in birds (García-Fernández et al. 2015; Kuenzel et al. 2015), and coordination of coiling behavior in developing zebrafish embryos (Friedmann et al. 2015). Heterologously expressed VA opsins from *Danio rerio* and *Xenopus tropicalis* showed an absorption maximum at ∼500 nm followed by a *cis-trans* isomerization of the retinal chromophore, forming a non-reversible active state that enables activation of G_i_ and G_t_ G proteins (Sato et al. 2011). Electrophysiological Patch Clamp measurements also confirm a G_i_-dependent, PTX-sensitive induction of potassium currents via opening of GIRK channels (Friedmann et al. 2015; Winans et al. 2023). Despite these characteristics, VAL opsins have not yet been applied for optogenetic applications.

Here we show that *_Ak_*VAL activates the full range of G_i/o_ family G proteins, while exhibiting fast kinetics, high sensitivity to low-intensity stimulation, and bistable properties in HEK293 cells and in purified protein samples. Furthermore, *_Ak_VAL* displays functional, bidirectional modulation of Purkinje cell firing patterns in cerebellar slices and allows for strong and robust effects on movement in C*aenorhabditis elegans* (*C. elegans*) *in vivo*.

## Results

### A VAL opsin from the retina of flashlight fish *Anomalops katoptron*

The nocturnal splitfin flashlight fish *Anomalops katoptron* (*A. katoptron*) lives in the Indo-Pacific region and can be found at the water surface in schools up to several hundred specimens during dark, moonless nights (Hellinger et al. 2017; Mark et al. 2018; Jägers et al. 2021; Jägers et al. 2024). *A. katoptron* is characterized by bioluminescent light organs, located under the eyes (Fig. 1a). Recently, we showed that *A. katoptron* expresses two different rhodopsins (Rh1and Rh2) in the retina, which are highly light-sensitive to detect blue bioluminescent light (Mark et al. 2018). Further, mRNA sequencing of *A. katoptron* retina specimen revealed an additional fulllength sequence of a vertebrate ancient long opsin homolog (*_Ak_*VAL). Phylogenetic analysis of *_Ak-_*VAL indicated a 94% sequence identity with the *Myripristis murdjan* (soldierfish of the Indo-Pacific) VAL opsin, and 76% identity to the VAL-a isoform of *Danio rerio* (zebrafish) (Fig. 1b; c). *_Ak_*VAL has a predicted typical seven transmembrane domains (7TM) with a conserved Schiff base lysine at TM7 (K308^7.10^) (Fig. 1c). Similar to other VAL opsins, *_Ak_*VAL also has the hallmark counterion at TM3 (E114 *^Ak^*^VAL^; E113 in bovine rhodopsin numbering), where the conserved glutamic acid shared between vertebrate and invertebrate opsins in the second extracellular loop (E181^ECL2^ in bovine) is substituted with a serine (S182^ECL2^; Fig. 1c) (Sato et al. 2011).

**Figure 1.**
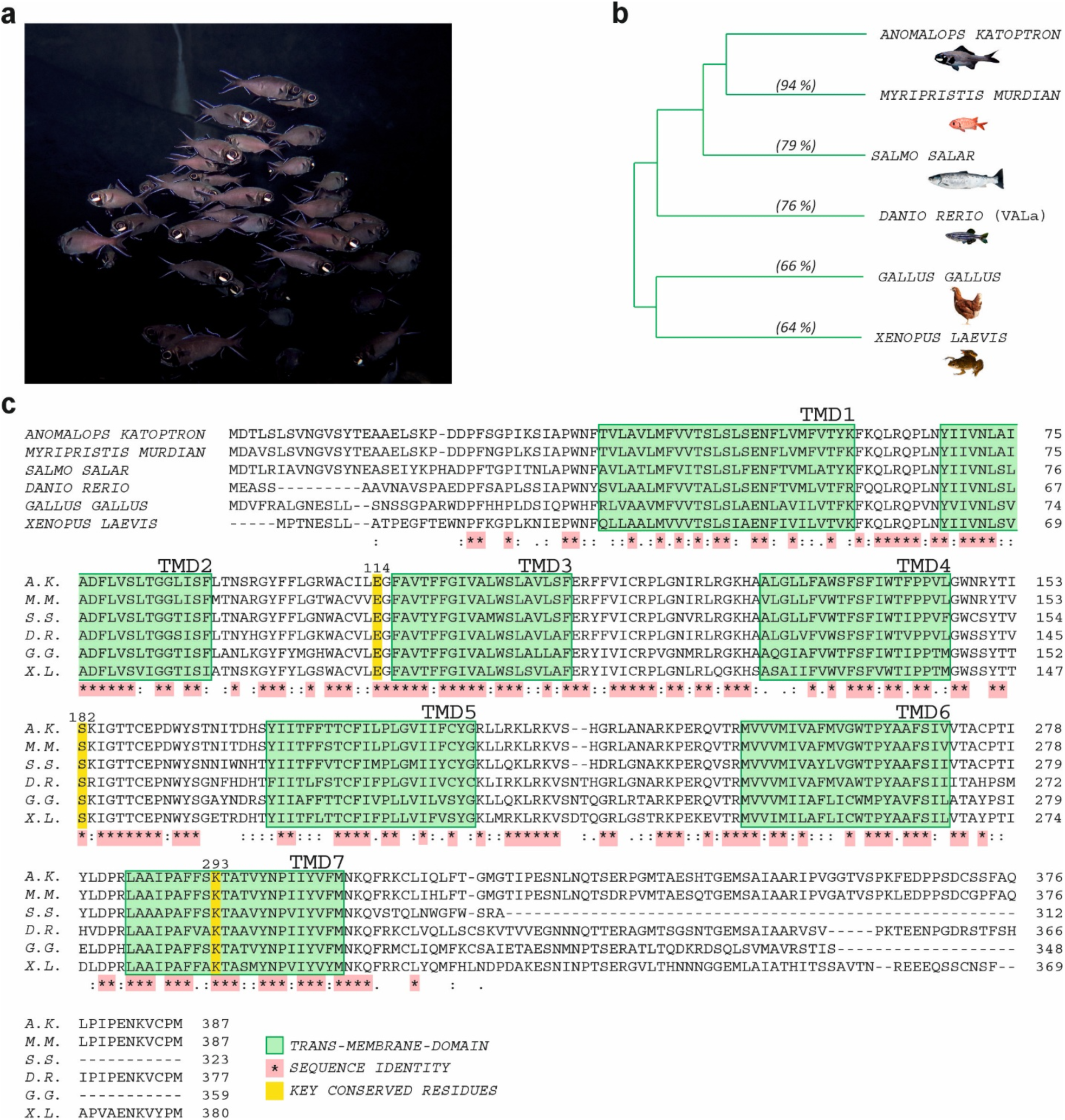
Phylogenetic analysis and structural properties of *Anomalops katoptron* VAL opsin. **a** Photograph of a group of *Anomalops katoptron* flashlight (photo credit: Stefan Herlitze) **b** Phylogenetic tree and sequence identity (%) of various VA opsins compared to _Ak_VAL. **c** Phylogenetic comparison of amino acid sequences of the vertebrate ancient long opsins across distinct species; green boxes indicate transmembrane-domains, key conserved positions in opsins including the classical counter ions E114 (113 in bovine rhodopsin (BR) numbering) and S182 (E181 in BR) and Schiff base lysine, K293 (K296 in BR) are highlighted in yellow. Notably, E181 is among the most conserved residues across all rhodopsin species is substituted with a serine residue (S182 in *_AK_*VAL) that is highly conserved amongst the VAL opsin family.

### *_Ak_*VAL is a bistable, G_i/o_ coupled light-activated GPCR

Having isolated the cDNA of *_Ak_*VAL, we first characterized the light-induced changes of the absorption spectra of a recombinant *_Ak_*VAL that was heterologously expressed in insect cells (Fig. 2, Supp. 1). Measurements of the absorbance of a *9-cis* retinal reconstituted *_Ak_*VAL indicated maximal absorbance at 490 nm. Following green illumination (520 nm), the pigment absorbance at ∼500 nm was significantly decreased while an increase was noted in the absorbance at the UV region (380 nm, Fig. 2). UV light irradiation converted *_Ak_*VAL back to a visible light-absorbing state with a λ max of approximately 500 nm. The conversion of UV to green and green to UV absorbing states could be induced repetitively with minimal decline in absorption peak amplitudes (Fig. 2b). A similar switchable behavior is also observed after 10 min dark incubation following the respective illumination at each specific wavelength (Supp. 1). The latter indicates that beyond having a biswitchable nature, _Ak_VAL is also thermostable. Notably, the maximal absorbance recorded for *_Ak_*VAL in the dark are in accordance with previous reports indicating that most VAL opsins are also sensitive to green light (e.g., zebrafish VAL-a and VAL-b absorb light at 510 nm and 505 nm respectively (Kojima et al. 2000; Kojima et al. 2008) and the western clawed frog’s VAL (*Xenopus tropicalis*) has a λ_max_ of 501 nm (Sato et al. 2011)). However, the reverse reaction has not been described for this opsin, and VAL opsins were thus considered to be non-bleaching, yet non-reversible type opsins (Sato et al. 2011). Such properties for an animal opsin were considered unique to the VAL opsin family and were attributed to the substitution of the conserved counter ion position, E181 (in bovine rhodopsin numbering), with a serine residue in the VAL opsin family (S182 in *_Ak_*VAL; Fig. 1c). Our results support a different photocycle for *_Ak_*VAL, that is indeed a non-bleaching opsin, but can be repetitively switched between two stable states (bistable), upon green and UV illumination cycles.

**Figure 2.**
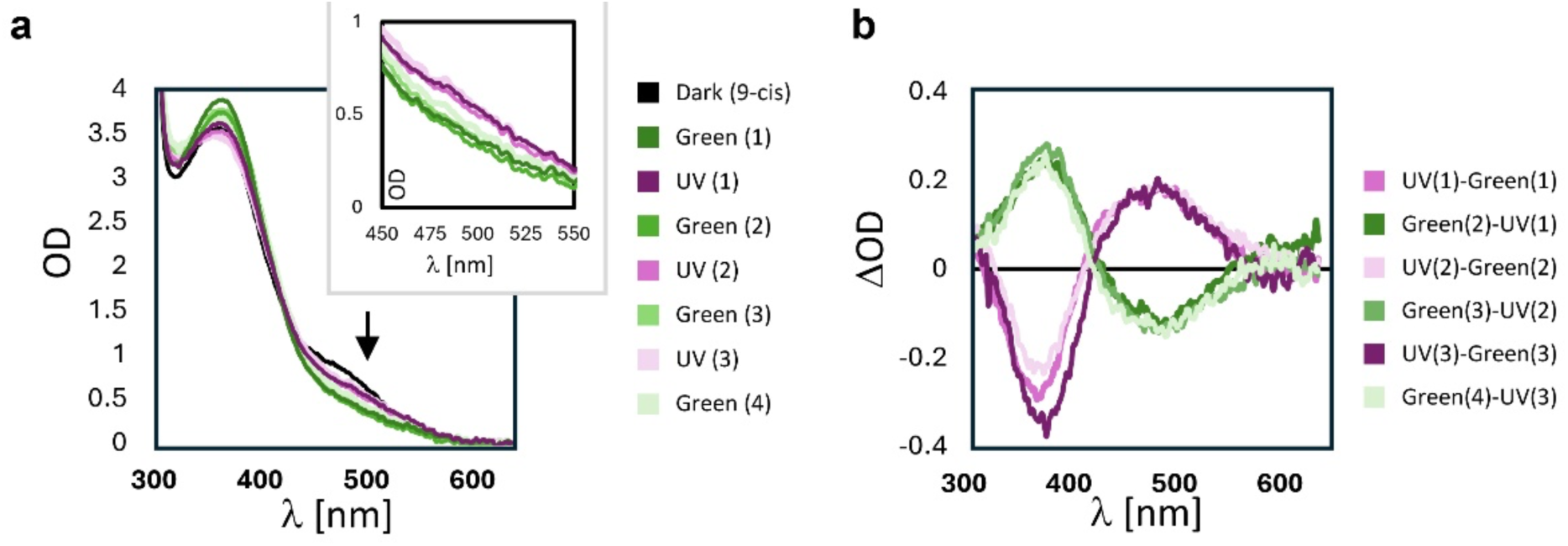
*_Ak_*VAL is a bistable receptor and reversibly responds to UV and green light. **a** Absorbance spectra of *_Ak_*VAL purified in detergents taken under differential illumination conditions. Plots are colored according to the index to the right with the dark state (black) containing *9-cis* retinal, followed by green (520 nm) and UV (390-400 nm) illuminations. Green illuminations are in different shades of green, UV illuminations are purple. Numbers in parenthesis correspond to the cycle number of illuminations (e.g., 1 corresponds to the first green illumination after dark etc.). The arrow indicates the absorbance peak at 495 nm (*11-cis*) that is slightly red shifted compared with the dark state sample (*9-cis*, black). The box in the right corner is a magnification of the 495 nm peak, showing that upon green illumination the OD is reduced, whereas after UV illumination the pigment converts back to the *11-cis* state. **b** Difference spectrum of the measurements presented in (a).

To further explore *_Ak_*VAL function, we expressed the cDNA encoding the full-length protein in HEK293 tsA cells and conducted signalling assays to evaluate G protein binding properties. To analyze the G protein binding selectivity, we performed the luminescent-based G protein activation assay (GsX-Assay) (Ballister et al. 2018) (Fig. 3a). In this experiment, the C-terminal alpha helix of the Gα_s_ subunit is substituted with the corresponding helix of a subset of Gα proteins, where cAMP production due to the activation of adenylyl-cyclase is used as a bioluminescence output; as the C-terminal domain is the main constituent of receptor binding, this assay allows for the rapid and high-throughput screen of binding partners. The experiments revealed that after blue light illumination, predominantly G_i_-family G protein subtypes are activated with the strongest luminescent increase of the G_t_ (5,24-fold) followed by G_z_ (4,91-fold), G_o_ (4,42-fold), and G_i_ (2,74-fold) proteins while no activation of G_s_- or G_q_-type proteins was observed (Fig. 3b). These results were confirmed by BRET assays following the dissociation of Gβγ from the Gα subunits (TRUPATH (Olsen et al. 2020)), that also indicated G_i_ and G_o_ but not G_q_-type signaling (Supp. 2). Furthermore, 2-Photon Calcium imaging showed no detectable changes in the G_q_-mediated intracellular Ca^2+^ levels (Supp. 3). In addition, GloSensor based cAMP assays using the endogenous G proteins expressed in HEK293 cells showed a reduction of endogenous cAMP levels upon light stimulation (Supp. 4). These results also support a G_i_-family subtype effect and are consistent with earlier reports showing that VAL opsin from *Xenopus tropicalis* and *Danio rerio* couples to G_i_ and G_t_ to induce G protein signaling (Sato et al. 2011).

**Figure 3.**
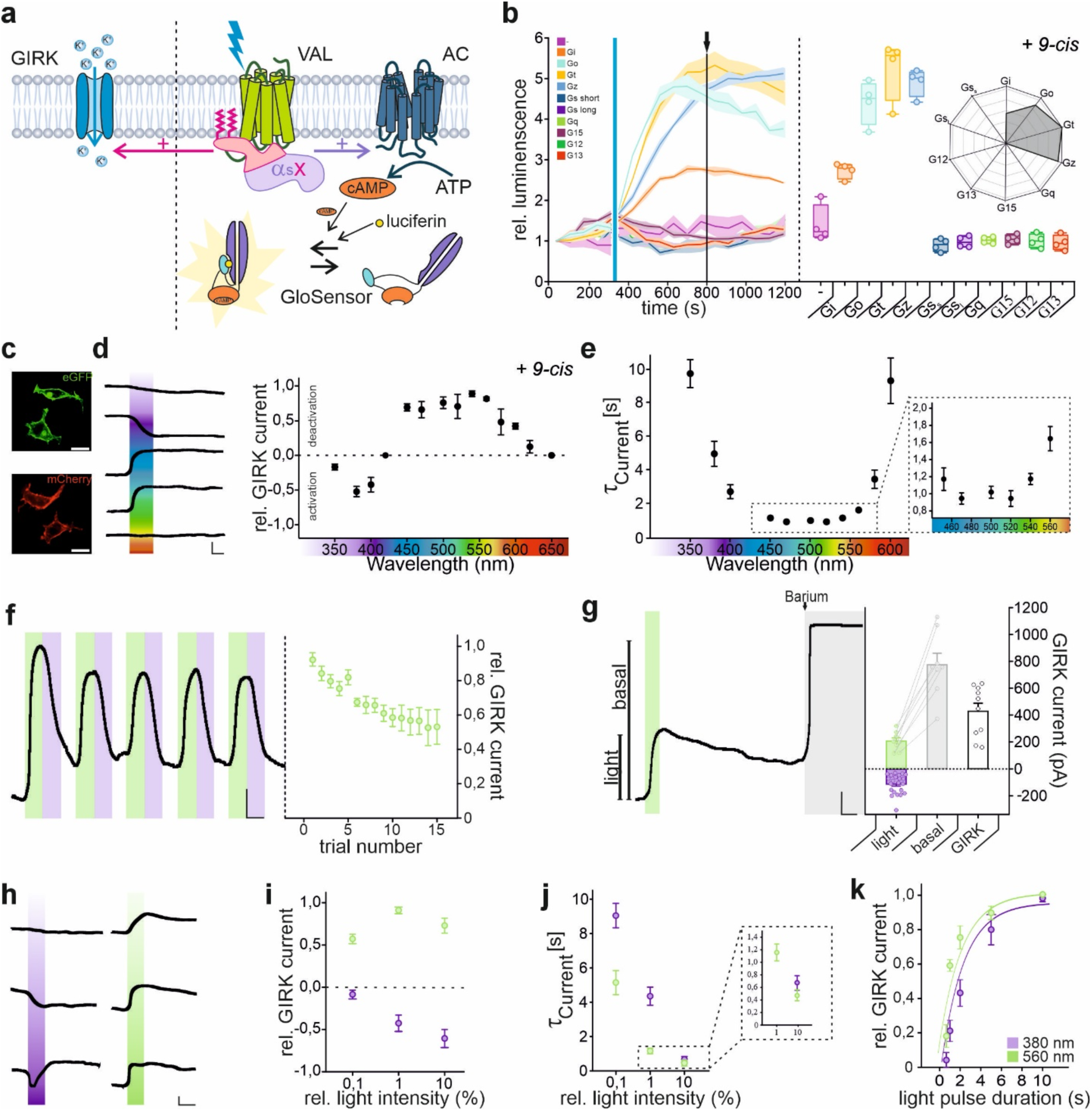
GsX Assay and electrophysiological characterization of photostimulated _Ak_VAL. **a** GIRK Assay (left) and GsX Assay (right) schematic overviews. In GIRK 1/2 expressing HEK cells dissociated G_i_ βγ-subunits activate GIRK channels and enable strong inward potassium currents (left). Gα binding domains of various G proteins are attached to the Gαs protein leading to a uniform cellular response (cAMP ↑) that can be measured via the GloSensor Assay in HEK tsA cells (right). **b** Relative luminescent traces for all tested G protein chimeras exhibit exclusive increase of luminescence after activation of _Ak_VAL with a 10 s blue light pulse only when coupled to the G_i_ family chimeras G_s_t, G_s_o, G_s_z and G_s_i (left); quantitative boxplot analysis and radar plot displaying relative luminescent increase (1x-fold each grid) of normalized luminescent values 800 s (black arrow) after the start of the measurement (right). **c** Expression of _Ak_VAL-eGFP (green, top) and _Ak_VAL-mCherry (red, bottom) in HEK293 tsA cells depicts predominant membrane expression; scale bar = 5 µm. **d-j** Whole-cell Patch Clamp recordings of _Ak_VAL-eGFP expressing HEK GIRK 1/2 cells supplied with 1 µM *9-cis* Retinal show G protein mediated GIRK-dependent potassium currents after low intensity irradiation. All recordings ranging from 420 nm upwards were performed after activation with 380 nm and all recordings below 420 nm after deactivation with 560 nm, mean values + standard errors are plotted in all cases. **d** Action spectrum of _Ak_VAL mediated photocurrents reveals a bistable like nature by inducing inward currents in the range of 350-420 nm and outward currents between 420-650 nm (n=15 cells) (right), example traces of a single cell illustrating the ion flow direction for 350,380,450,560 and 650 nm (left); scale bar: 50 pA/. **e** _Ak_VAL exhibits extraordinarily fast sub second off kinetics in the range of 450-540 nm (zoomed in panel, right) and slightly slower on kinetics (left) (n=15 cells). **f** Example trace depicting _Ak_VAL to be repeatedly de-/activatable switching between 380 and 560 nm light stimuli (left) while lowering the deactivation amplitude to ∼ 50 % within 15 repetitions (right); scale bar: 50 pA/10 s. **g** Representative trace indicates basal activity of _Ak_VAL supplied with 1 µM *9-cis* retinal in the dark state (left) that exceeds the GIRK leak current (white bar) of untransfected cells. While the VAL induced basal current can be temporarily reduced by green light irradiation (560 nm) it is abolished completely after application of 5 mM barium chloride solution (black arrow, grey area); scale bar: 100 pA/10 s; quantitative comparison of light induced current compared to basal current (n=8 cells) (right). **h-j** Analysis of intensity dependance of _Ak_VAL mediated GIRK currents displays outstanding sensitivity and kinetics. **h** GIRK current de-/activation maximizes between 0,1-100 µW/cm^2^ 560/380 nm stimulation; scale bar: 100 pA/10 s. **i** Quantitative analysis of intensity-dependent evoked currents (n=9 cells each) (right). **j** Intensity based kinetics measured between 0,1-100 µW/cm^2^ for 380 and 560 nm (left) reveal sub second on and off kinetics (zoomed in panel, right) (n=9 cells each). **k** Comparison of different light stimuli periods to de-/activate _Ak_VAL opsin, GIRK currents saturate in both cases from durations reaching 5 s and can still be measured down to 0,5 s (n=6 cells each).

To gain a deeper insight into the action spectra and biophysical/physiological properties of *_Ak_*VAL, we focused on the G_i_ pathway activation and characterized light-induced, G_i_ βγ-subunit-dependent G protein-gated inwardly rectifying potassium (GIRK) currents using Patch Clamp recordings. In the presence of the *9-cis* retinal *_Ak_*VAL-eGFP/mCherry expressing HEK GIRK 1/2 cells (Fig. 3c) showed an increase in baseline current by stimulation with UV/blue light (360-400 nm) and a reduction in baseline current in response to blue-red light (420-620 nm) stimulation with a λ_max_ of ∼550 nm – green (Fig. 3d). This indicates an _Ak_VAL-dependent activation of GIRK channels by UV light and a deactivation of GIRK channels by green light which reflects the bistable characteristics observed *in vitro*. *_Ak_*VAL exhibited fast sub-second off kinetics during blue-green light deactivation and slower activation kinetics of 2,73 ± 0,39 s at 400 nm irradiation (Fig. 3e). The activation and deactivation of the GIRK current could repetitively be induced with a decline in response amplitude by around 50 % after 15 trials (Fig. 3f). Light-induced GIRK currents were inhibited by the addition of the G_i_ blocker PTX but remained unaffected after adding the PLC blocker U73122 which shuts down G_q_-dependent Calcium release within the cells (Supp. 5). To investigate the contribution of constitutive dark activity of GIRK currents by the expression of *_Ak_*VAL, we blocked the GIRK current after light activation with Barium and compared the light-modulated (green light) currents to the baseline current. The results suggest that ∼20-30 % of the basal GIRK current can be increased or reduced by UV/blue or green light, respectively (Fig. 3g).

We next analyzed light intensity-dependence of activation and deactivation of *_Ak_*VAL induced GIRK currents (Fig. 3h-j). GIRK current activation increases with increasing UV light and deactivation increases with increasing green light intensities. Green light of 3-5 µW/cm^2^ intensity already induced a 50% deactivation of the GIRK current (Fig. 3h-i). Using low light conditions, we also determined the *_Ak_*VAL-mediated, wavelength-dependent activation/deactivation times. The activation and deactivation times also depended on the relative light intensities accelerating with higher light intensities of approximately τ= 0.7 sec for activation and 0.5 sec for deactivation. We found that the time of UV/blue (360-400 nm) light-dependent GIRK channel activation decreased from several seconds to sub-second activation, while blue/green (440-540 nm) light-dependent GIRK channel deactivation kinetics were very fast of around 1 sec (Fig. 3j; Zoom). The amplitude of both activated and deactivated GIRK currents also depended on the light-pulse duration, exponentially increasing with half maximal activation at around 2s duration (Fig. 3k).

To investigate if the anomalous activation spectra of *_Ak_*VAL are species-specific, we compared the chromophore-dependent action spectra and bistable properties of VAL opsin from *A. katoptron* with that of the VAL opsin analog from the zebrafish *Danio rerio* (76% AA sequence identity) (Kojima et al. 2000; Friedmann et al. 2015). As indicated in Supp. 6, VAL opsin from *Danio rerio* reveals the same chromophore-dependent action spectra and bistable properties as *_Ak_*VAL.

### *_Ak_*VAL bidirectionally modulates cerebellar Purkinje cell activity

According to the light-dependent modulation of GIRK channels by *_Ak_*VAL in HEK293 cells we hypothesized that UV light, which activates G_i/o_ signaling, may decrease, while green light stimulation, which deactivated G_i/o_ signaling, increases neuronal activity. We could previously show that light activation of opto-GPCRs can modulate spontaneous firing activity of cerebellar Purkinje cells (Gutierrez et al. 2011; Surdin et al. 2023; Mücher et al. 2025). Therefore, we expressed *_Ak_*VAL-eGFP in cerebellar Purkinje cells (Fig. 4a) and examined if light stimulation of *_Ak_*VAL modulates neuronal activity in a wavelength-specific manner. As demonstrated in Figure 4, 560 nm light stimulation increased (Fig. 4b-c), while 380 nm light reduced the spontaneous AP firing rate of *_Ak_*VAL expressing Purkinje cells (Fig. 4d-e). The light-dependent modulation of Purkinje cell excitability was further revealed using a current step protocol (Fig. 4f-g). As observed for the spontaneous AP firing rate, 560 nm light stimulation again increased the current step-induced spike rate (Fig. 4h) and led to a decrease in the spike threshold/rheobase (Fig. 4i). In contrast, 380 nm light stimulation decreased the induced spike rate (Fig. 4j). The robust changes in neuronal excitability by 560 nm light stimulation are consistent with a decrease in the membrane potential (Fig. 4k) and decrease in the latency to the first AP (Fig. 4l). 380 nm light stimulation did not show any significant changes in the rheobase, membrane potential or latency to the first action potential. However, the change in neuronal excitability induced by both light conditions recovered after the cell was kept in the dark for at least 5 min (Fig. 4m). In addition, an increase in input resistance was observed for 560 nm light stimulation using hyperpolarizing current steps (Fig. n-o), indicating a reduction in open ion channels in the cell membrane, e.g. GIRK channels (Fig. 4p).

**Figure 4.**
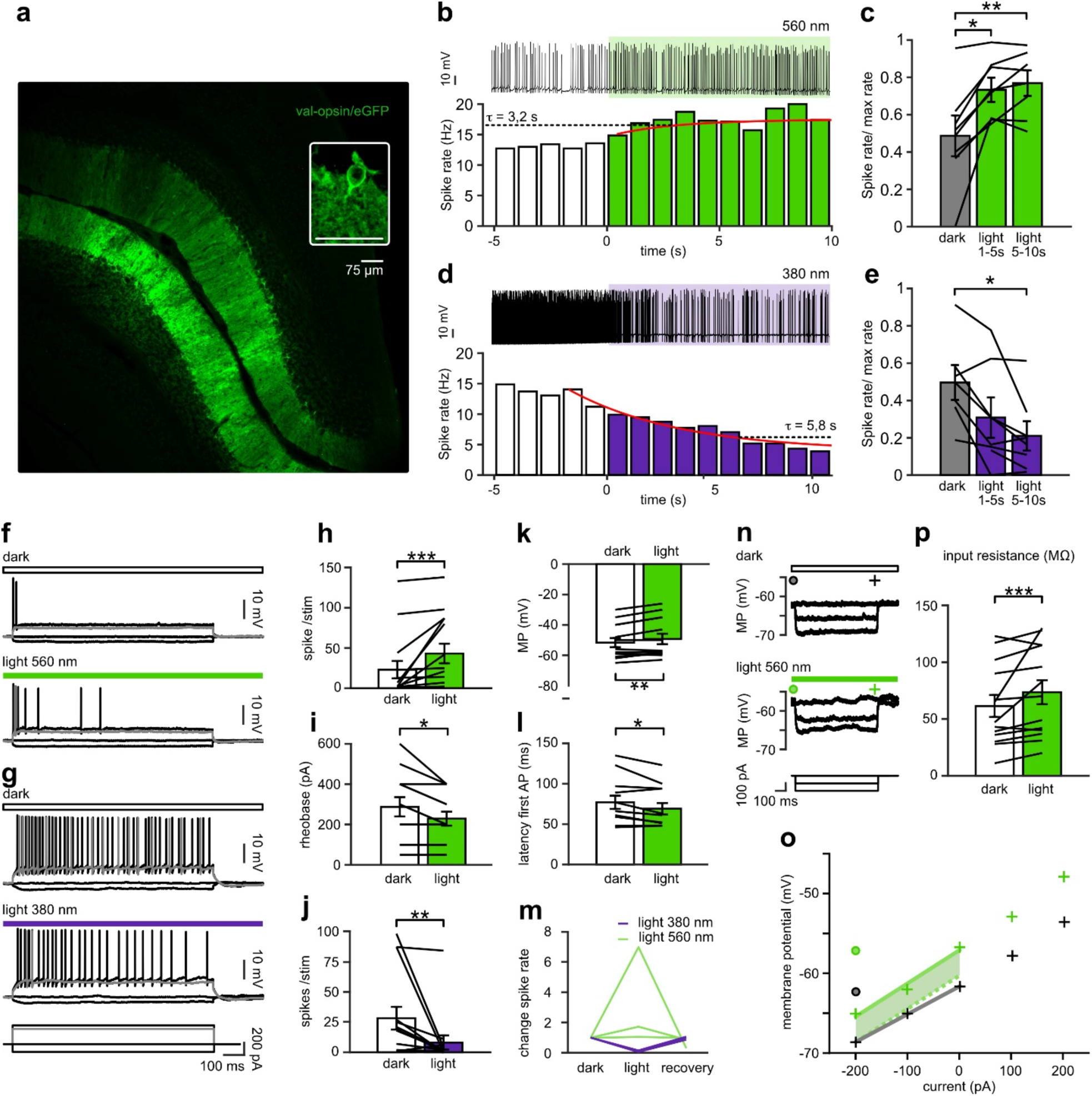
_Ak_VAL bidirectionally modulates PC firing in cerebellar slices. **a** Cerebellar slices showing VAL-GFP-expressing Purkinje cells. **b** On the top an example recording of a single neuron stimulated with a 10s-540 nm light (green area). Light stimulation increased the mean firing rate of all spontaneous firing neurons (bottom; n = 7). **c** Spike rate increase after 540 nm-light stimulation (F(2,28) = 17, p<0.0001; Significance was assessed using repeated measures ANOVA test followed by multi comparison, dark/light 0-5s: p = 0.016; dark/light 5-10s: p = 0.004). **d** On the top is an example recording of a single neuron stimulated with a 10s-380 nm light (purple area). Light stimulation decreased the mean firing rate of all spontaneous firing neurons (bottom; n = 7). **e** Spike rate decreases after 540 nm-light stimulation (F(2,28) = 17, p<0.0001; Significance was assessed using repeated measures ANOVA test followed by multi comparison, dark/light 0-5s: p = 0.016; dark/light 5-10s: p = 0.004). **f** Example neuronal response of single neurons expressing VAL to a −100, 0 300 and 400 pA pulses before stimulation (top) and during green light stimulation (bottom). **g** Example neuronal response of single neurons expressing VAL to a −100, 0 300 and 400 pA pulses before stimulation (top) and during purple light stimulation (bottom). **h** Mean response rate of VAL expressing cells increased during green light stimulation (n = 13; p = 0.0002). **i** Mean rheobase of VAL expressing cells decreased during green light stimulation (n = 11; p = 0.03). **j** Mean response rate of VAL expressing cells increased during purple light stimulation (n = 14; p = 0.002). **k** Mean membrane potential of VAL expressing cells increased during green light stimulation (n = 13; p = 0.002). **l** Mean latency of the first action potential (AP) of VAL expressing cells decreased during green light stimulation (n = 11; p = 0.04). **m** Input resistance membrane potential of VAL expressing cells increased during green light (n = 13; p = 0.0008). **n** Example response of a VAL expression neuron to hyperpolarizing current steps before and during light stimulation. **o** Input resistance was calculated using the slope of the membrane potential plotted against the hyperpolarizing current step. **p** Spike rate change recovered a recovering period of at least 5 minutes after purple light stimulation.

### *_Ak_*VAL strongly inhibits motor behavior in *C. elegans in vivo*

We next investigated if *_Ak_*VAL can be functionally expressed in a invertebrate model system (i.e. C. elegans), and whether a light-activation of *_Ak_*VAL would modulate G_i/o_-mediated behaviors. Expressing *_Ak_*VAL under the control of the *unc-17* promotor led to robust expression throughout the cholinergic nervous system including the nerve ring as well as the dorsal and ventral nerve cords in *C. elegans* (Fig. 5a). Unlike mammalian systems, *C. elegans* does not produce endogenous retinoids, so the supply of retinal is essential for opsin functionality (Fig. 5b-c). Blue light excitation of *9-cis* retinal-fed animals resulted in a severe, long-lasting reduction in crawling speed that persisted for minutes after the stimulus ended, while control animals, which did not receive *9-cis* retinal, showed no response to the light pulse (Fig 5b, Supp. Videos 1, 2). The *_Ak_*VAL mediated light-induced reduction in crawling speed is correlated with an increase in body length (Fig. 5c). In addition, we were able to confirm *_Ak_*VAL mediated bidirectional modulation of the G_o_ pathway in a light- and retinal-dependent manner (Fig. 5 e-f). Consistent to our biophysical, cell- and neurophysiological data we were able to inhibit the crawling speed of *_Ak_*VAL expressing individuals after green light stimulation and abolish the effect with subsequent UV light (Fig. 5e). Without subsequent UV excitation, *C. elegans* movement remained impaired for several minutes (Fig. 5f). The activation of *_Ak_*VAL in cholinergic neurons is supposed to reduce the acetylcholine release via the activation of the G_o_ pathway. *GOA-1* is the only G_i/o_ type G protein encoded in the genome of *C. elegans* and is likely activatable by *_Ak_*VAL due to an 80 % amino acid sequence similarity compared to mammalian G_o_ proteins. Thus, the increased body length in reaction to UV light is likely caused by the relaxation of the body wall due to the hyperpolarization of the muscle tone by the activation of G_o_ in the corresponding motor neurons (Fig.5c). To confirm this hypothesis, we generated a cross between *_Ak_*VAL expressing animals and the *GOA-1* knockout strain which lacks G_i/o_ type of G proteins. Compared to Wt *_Ak_*VAL expressing animals, the *GOA-1* knockout significantly reduces *C. elegans* response to light stimulation. This finding substantiates the hypothesis that *_Ak_*VAL opsin has the potential to activate an evolutionary conserved G_o_ pathway (Fig. 5d).

**Figure 5.**
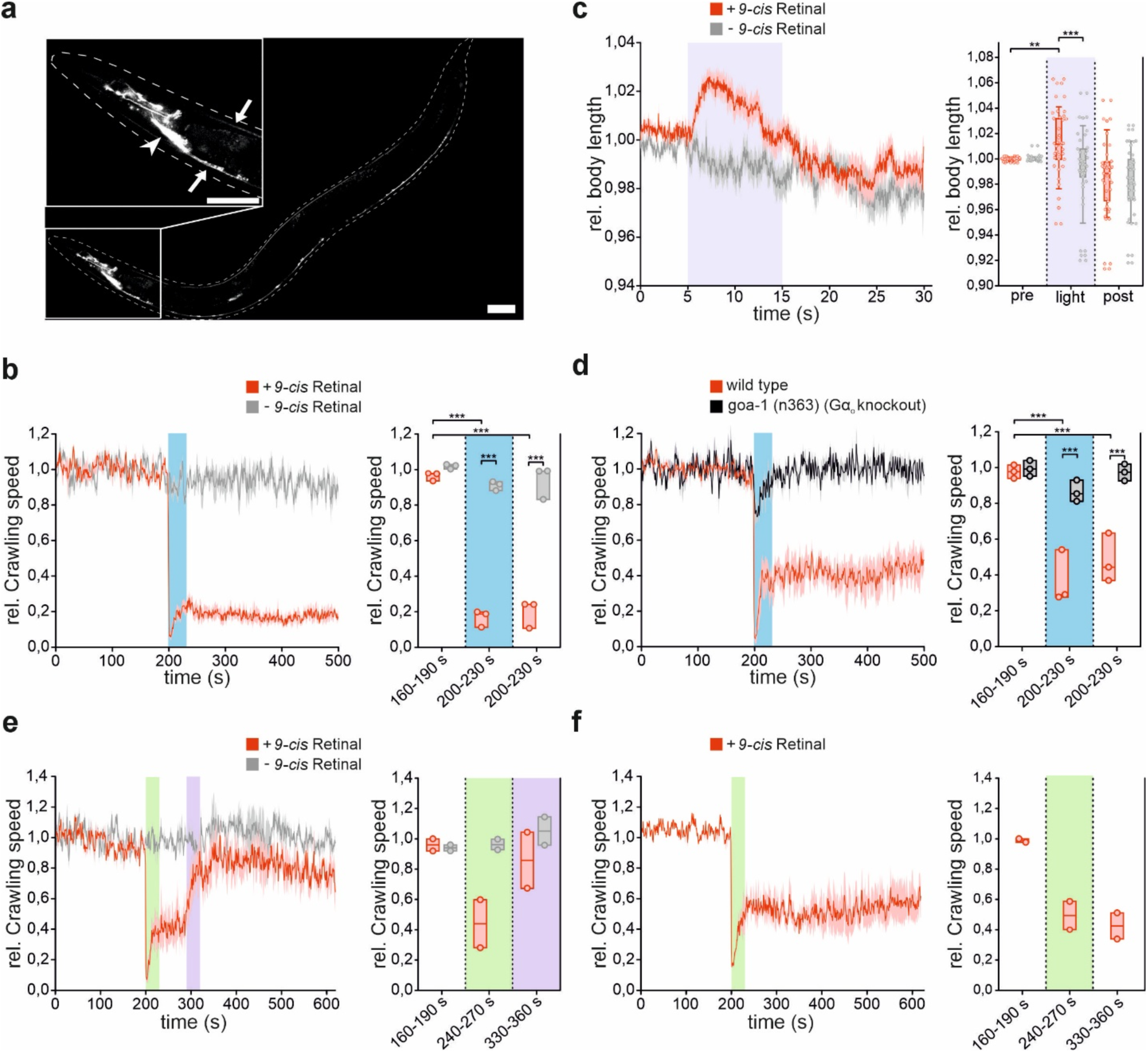
_Ak_VAL opsin inhibits motor behaviour in *C. elegans.* **a** Representative image of *_Ak_*VAL-mCherry expression in cholinergic neurons of *C. elegans* including close-up of the head ring region. Arrowhead indicates the nerve ring (central nervous system), arrows indicate dorsal (top) and ventral (bottom) nerve cords. The outline of the animal is marked by a dashed line; Scale bar = 50 µm. **b** Mean (± SEM) normalized crawling speed optionally treated with *9-cis* retinal. A 30 s light pulse (470 nm, 0.1 mW/mm^2^) was applied after 200 s as indicated by the blue bar (left); Statistical boxplot analysis of crawling speed data. Each dot represents a population of animals tested on a single day. Two-way ANOVA with Tukey’s correction for multiple comparisons. N = 3 populations were tested (right). **c** Mean (± SEM) normalized body length of animals expressing _Ak_VAL-mCherry in cholinergic neurons optionally treated with *9-cis* retinal. A 10 s light pulse (0.1 mW/mm^2^, 380 nm) was applied after 5 s as indicated by purple shade. Number of animals measured (n) accumulated from N = 3 biological replicates (left), Statistical boxplot analysis of body length data. Each dot represents a single animal. Two-way ANOVA with Tukey’s correction for multiple comparisons (right). **d** Mean (± SEM) normalized crawling speed of animals expressing _Ak_VAL-mCherry in cholinergic neurons treated with *9-cis* retinal. Genotype of animals is indicated in legend. A 30 s light pulse (470 nm, 0.1 mW/mm^2^) was applied after 200 s as indicated by blue bar (left). Statistical boxplot analysis of crawling speed data. Each dot represents a population of animals tested on a single day. Two-way ANOVA with Tukey’s correction for multiple comparisons N = 3 populations were tested (right). ** p < 0.01, *** p < 0.001. **e-f** Mean (± SEM) normalized crawling speed optionally treated with *9-cis* retinal. A 30 s green light pulse (525 nm, 65 µW/mm2) was applied after 200 s as indicated by the green bar, optionally followed by a 30s UV light pulse (365 nm, 25 µW/mm2) after 60 s as indicated by a violet bar (left) (**f**); statistical boxplot analysis of crawling speed data. Each dot represents a population of animals tested on a single day. N= 2 populations were tested (right).

## Discussion

### Ancient long wavelength opsin in teleosts

Vertebrate ancient (VA) opsins belong with pinopsin, parapinopsin and parietopsin to the non-visual opsins in the vertebrate opsin subfamily and are more than 40% identical with the visual opsins (Shichida und Matsuyama 2009). These opsins have been suggested to be involved in regulation of circadian rhythms, photoperiodicity and body color change. VA opsins have been first identified in salmon (Soni und Foster 1997). Two VAL opsin variants (a and b) are found in teleosts and represent elongated isoforms of VA opsins. They are characterized by their extension of their C-terminus and their differential expression suggesting different physiological roles (Kojima et al. 2000; Kojima et al. 2008). VAL opsins are expressed throughout the body but in particular in the retina (horizontal cells) (Jenkins et al. 2003; Kojima et al. 2000) brain, testis and skin, where the function of VAL opsins have only been addressed rudimentarily. Knockout studies and observations in zebrafish suggest that VAL opsins play a role in chorion formation and embryonic hatching (Hang et al. 2016). In addition, isoform A of VAL opsin is expressed in the cilia of spinal neurons early in development and most likely mediates the inhibition of locomotion by environmental light via G_i/o_-mediated inhibition of the spinal central pattern generator (Friedmann et al. 2015; Winans et al. 2023). In our studies we identified one VAL opsin variant (homologue to isoform A) in the retina, brain and skin of *A. katoptron* using mRNA sequencing, where a physiological role has not been described so far.

### Ancient long wavelength opsin of *A. katoptron* is a bistable opsin

Functional reconstitution of VAL opsin from *Xenopus tropicalis* and *Danio rerio*, but not of VA opsin, in HEK293 cells with *11-cis* retinal and analysis of the absorption spectra revealed absorption and activation peaks around 500 nm (501 and 505 nm respectively, green absorbing pigment) (Kojima et al. 2000; Friedmann et al. 2015; Winans et al. 2023). The photoreaction is accompanied by *11-cis* to *all-trans* isomerization of the retinal chromophore and a shift in the absorption spectra from 501 to 455 nm when irradiated with >500 nm light (Sato et al. 2011). Subsequent irradiation of the photoproduct with 439 or <400 nm light did not cause an absorbance change suggesting that VAL from *Xenopus tropicalis* is not a bistable pigment. Nevertheless, while the photoproduct could not be isomerized back through illumination, these studies indicated that the retinal does not bleach, thus the opsin has been defined as a non-bleaching, non-bistable specie. In contrast, our studies show that in purified *_Ak_*VAL that has been reconstituted with *9-cis* retinal, repeated >500 nm (green) and <400 nm (UV) illumination cycles lead to a reversible photoconversion between two thermally stable retinal pigment states that most likely correspond to *all-trans* and *11-cis*. Bistability for _Ak_VAL was further demonstrated *in vitro* in HEK cells and in neurons, and through *in vivo* studies in *C. elegans*.

*_Ak_*VAL activates specifically G proteins of the inhibitory G protein family (G_i_, G_o_, G_t_, and G_z_), with no activation of the G_q_ and G_s_ pathway. Here we now confirmed this specificity in Gi/o coupling using various *in vitro* assays in HEK cells and an *in vivo* approach in *C. elegans* using a knock-out strain. Therefore, *_Ak_*VAL is suited to control the G_i/o_ signaling pathways in neuronal circuits and other cell-types. Various light-activated GPCRs have been adapted to control the G_i/o_ pathway including vertebrate rod and cone opsins (Li et al. 2005; Oh et al. 2010; Masseck et al. 2014; Tichy et al. 2019), lamprey parapinopsin (Eickelbeck et al. 2020; Copits et al. 2021), mosquito rhodopsin (Mahn et al. 2021) and recently various other bistable opsins including *Platynereis dumerilii* ciliary opsin (*Pd*CO) (Wietek et al. 2024). *_Ak_*VAL expands the optogenetic toolbox as a comparably very sensitive receptor characterized by its fast and red-shifted off properties, G_i_ pathway specificity, and bistability. Surprisingly, bistable opsins have not been used so far to control neuronal activity bidirectionally *in vitro* and *in vivo*. We explored this possibility in cerebellar Purkinje cells. In HEK293 cells expression of *_Ak_*VAL increases the constitutive activity of GIRK channels by 81 %. This allows for increasing and decreasing GIRK channel activity by UV/blue light and green/red light, respectively. Therefore, in excitable cells increasing the activity of the G_i/o_ pathway by UV/blue light should decrease excitability, while decreasing the G_i/o_ activity by green/red light should increase neuronal excitability. Indeed, 380 nm light pulses decreased, while 540 nm light pulses increased the spiking activity of *_Ak_*VAL expressing Purkinje cells. We also explored the possibility to use *_Ak_*VAL in an invertebrate animal model system. Expression of *_Ak_*VAL in cholinergic neurons of *C. elegans* induced strong, bidirectional G_o_-dependent behavioral effects, i.e. reduction in crawling speed correlated with an increase in body length. The behavioral effects are dependent on the addition of *9-cis* retinal and can be switched on and off by blue/green and UV/blue light, respectively. Thus, *_Ak_*VAL can be used to robustly induce and bidirectionally control G_i/o_ activity by UV/blue and green light.

Besides *_Ak_*VAL two other UV-activated, bistable vertebrate opsins have been described so far, i.e. parapinopsin (Blackshaw und Snyder 1997) and Opn5 (Tarttelin et al. 2003; Tomonari et al. 2008). All three opsins are UV-sensitive in the *9/11-cis*-retinal bound state, which converts the opsins to a G protein activated, *all-trans*-bound state. The active state can be converted to the resting state by blue-green light. The homology between *_Ak_*VAL and Opn5m and parapinopsin and Opn5m is low (28% and 26%) but is around 44% between *_Ak_*VAL and parapinopsin. These differences in homology may explain the differences in G protein specificity. *_Ak_*VAL and parapinopsin specifically activate the G_i/o_ pathway, while Opn5m activates G_i/o_- and G_q_-type G proteins (Sato et al. 2023; Wagdi et al. 2022) suggesting differences in structural mechanisms of G protein activation. Thus, it will be interesting to investigate the common features and structural difference of the different bistable, UV-sensitive opsins.

In summary, we identified a new bistable, G_i/o_ coupled opsin variant *_Ak_*VAL, for light-dependent bidirectional control of G_i/o_ pathways in neuronal networks from mice and powerful G_o_-mediated modulation of the neuronal networks of *C.elegans*.

## Supplementary

**Supplementary Figure 1:**
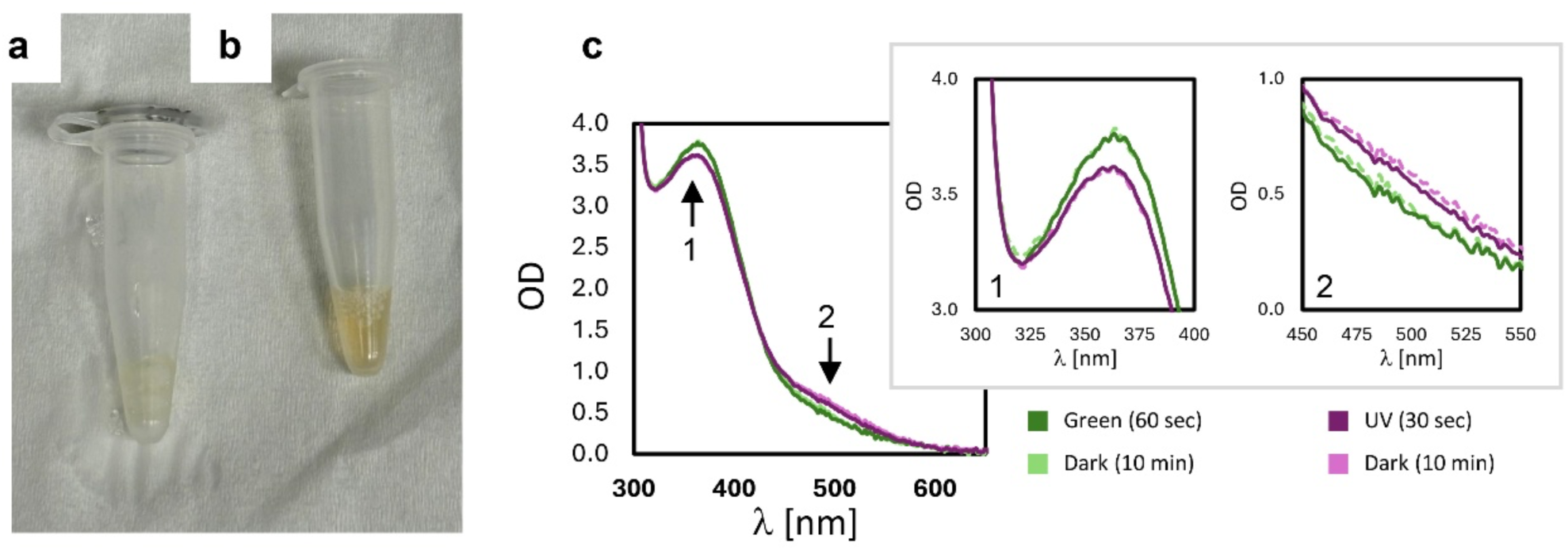
*_Ak_*VAL produces a thermostable active state upon illumination. **a-b** The color of the *_Ak_*VAL protein sample purified in detergents after illumination with green (a) and UV (b) lights. **c** Absorbance spectra of *_Ak_*VAL purified in detergents taken under differential illumination conditions. Plots are colored according to the key presented below the figure. Arrows indicate the peaks corresponding to the *all-trans* (1) and *11-cis* (2) retinal configurations. Peaks are enlarged in the right top box, to show that following a 10 minutes dark incubation after each illumination cycle, the sample was thermostable, and did not revert fully or partially to the reciprocal pigment state.

**Supplementary Figure 2:**
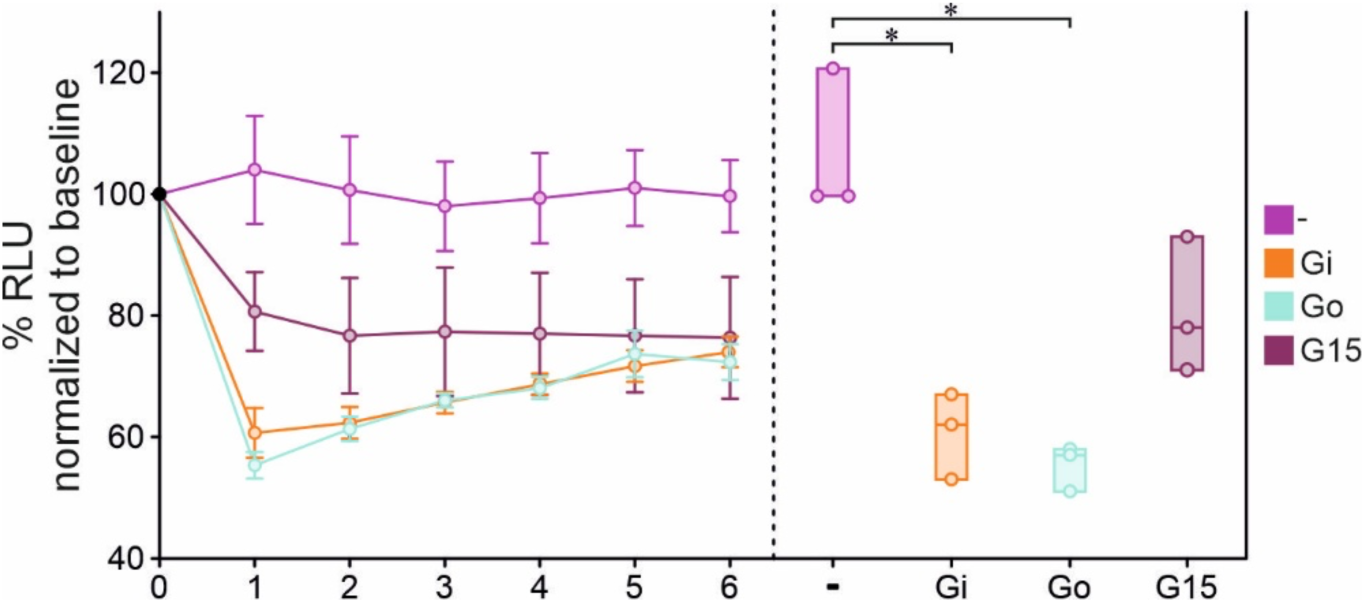
TRUPATH Assay demonstrates predominant activation of G_i_ and G_o_ over G_q/15_ type G proteins. **a** G protein coupling activity of *_Ak_*VAL after activation with light at 385 nm for 1 s in HEK293ADG7 cells show specific G_i_ and G_o_ coupling. Data are % BRET ratio (410/515 nm) normalized to baseline responses and are shown as mean ± SEM (left); statistical boxplot analysis of relative BRET ratio decrease (%) at the first measurement after illumination. Each dot represents one independent experiment (N=3); Kruskal-Wallis One Way Analysis of Variance on Ranks performed for statistical significance, *=p<0,05.

**Supplementary Figure 3:**
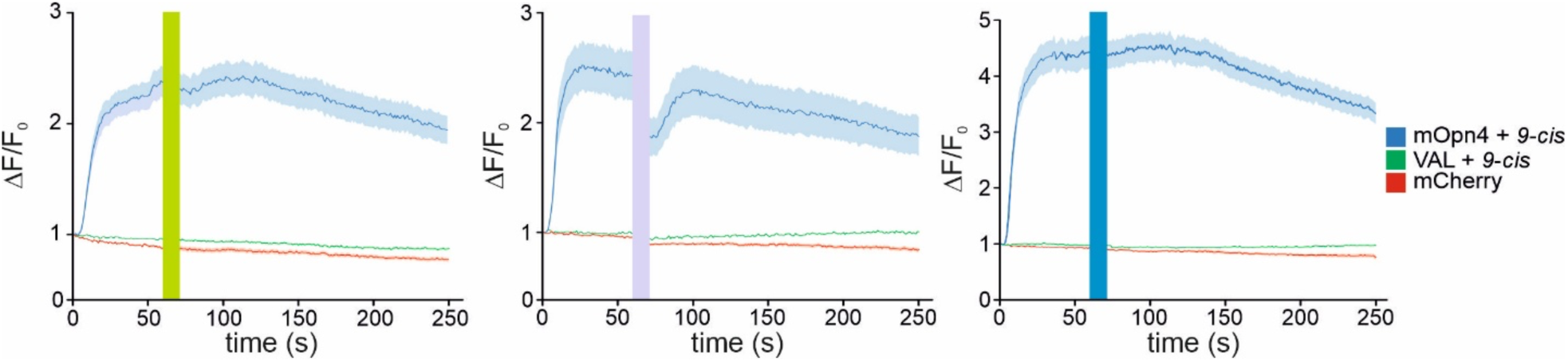
_Ak_VAL opsin does not influence intracellular Calcium dynamics upon activation. **a** 2-Photon Calcium Imaging of HEK293 tsA cells coexpressing *_Ak_*VAL and GCaMP6m activated by 560 nm (left), 380 nm (middle) or 480 nm (right) did not result in changes of fluorescence while the G_q_-coupled mOpn4L elevates intracellular Calcium concentrations. Cell culture dishes are constantly illuminated with 965 nm 2P-laser to monitor GCaMP fluorescence, (n=3 wells; 80-200 cells, line=mean, shaded area=SEM).

**Supplementary Figure 4:**
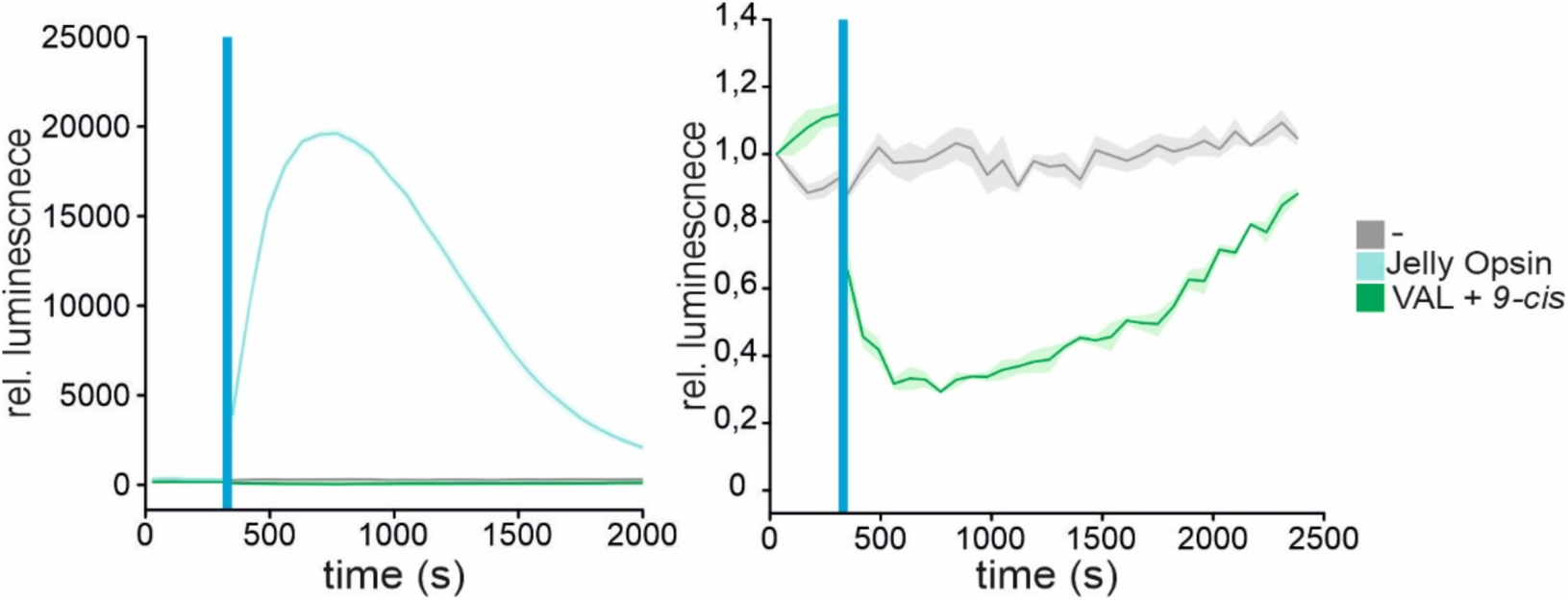
_Ak_VAL opsin does not increase but reduces endogenous cAMP levels. Luminescence based GloSensor Assay to monitor intracellular cAMP (n=4 wells, line=mean, shaded area=SEM). Normalized luminescent values display Jelly Opsin robustly induces strong increase of cAMP concentration upon activation with 480 nm blue light while *_Ak_*VAL does not (left) but rather reduces endogenous cAMP to ∼ 30 % of baseline levels (right).

**Supplementary Figure 5:**
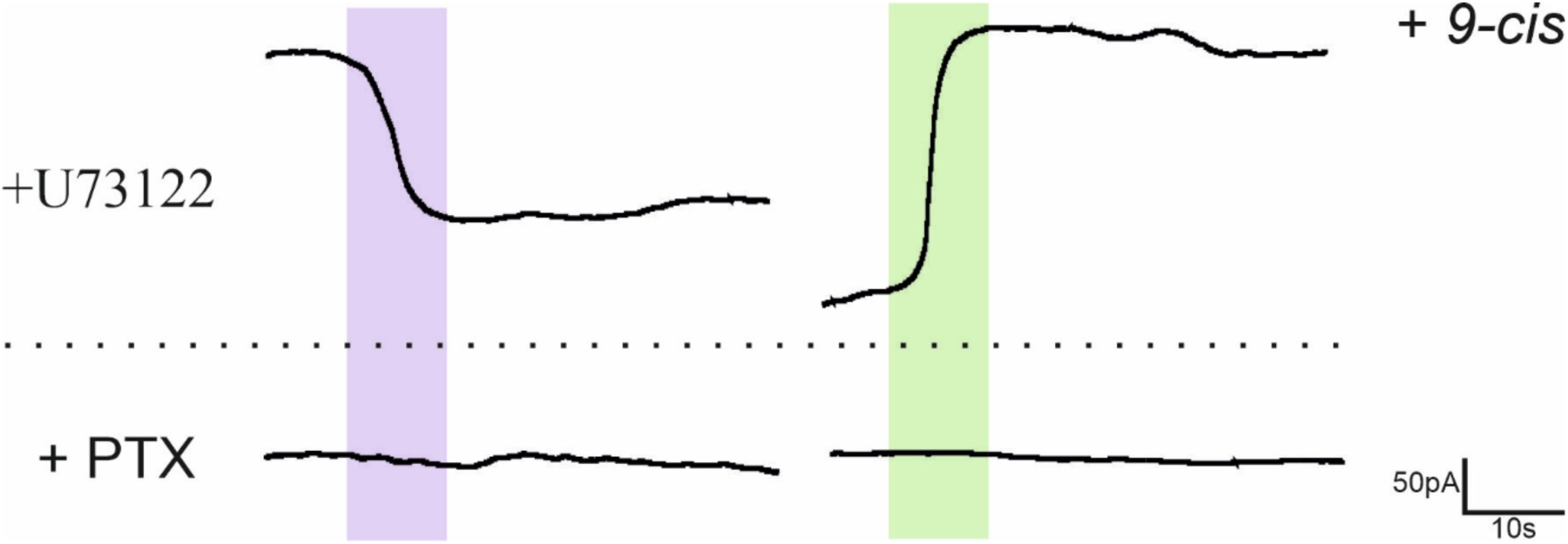
Inhibition of G_i_ proteins but not PLC activity blocks _Ak_VAL induced GIRK currents. Whole-cell Patch Clamp recordings of 1 µM *9-cis* Retinal supplied HEK GIRK 1/2 cells stimulated with 380 nm UV or 560 nm green light. Addition of 10 µM of the PLC inhibitor U73122 did not affect GIRK currents while 24 h incubation with 200 ng/ml of the G_i_ protein inhibitor PTX completely negates all currents.

**Supplementary Figure 6:**
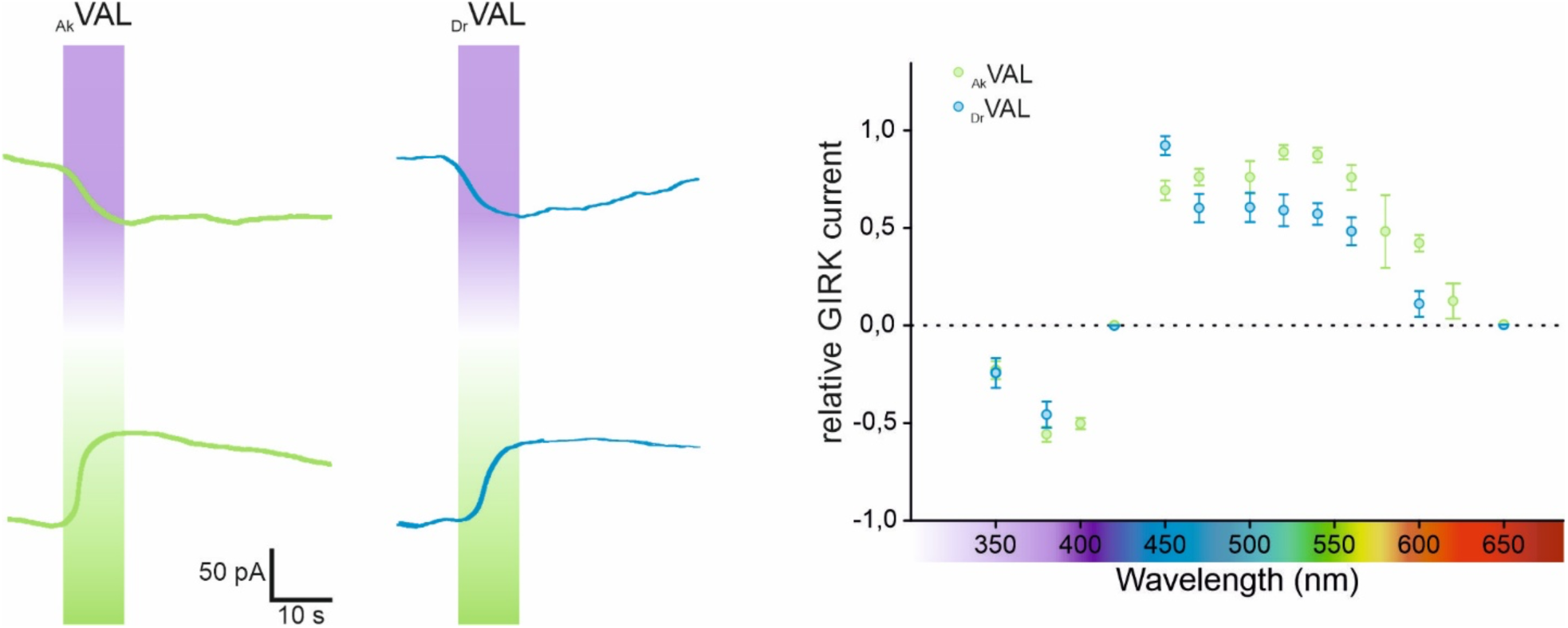
Electrophysiological comparison of *Anomalops katoptron* VAL and *Danio rerio* VAL: Whole-cell Patch Clamp recordings of *_Ak_*VAL-eGFP and *Danio rerio* VAL (*_Dr_VAL*) expressing HEK GIRK 1/2 cells supplied with 1 µM *9-cis* Retinal show no difference in bistable properties when illuminated with low intensity UV/green light (left) and no spectral differences in G protein mediated GIRK-dependent potassium currents after low intensity irradiation (right).

## Methods

### Transcript analysis and sequence alignments

*Anomalops katoptron* retinal mRNA samples were prepared to synthesize VA opsin cDNA using *the SuperScript III First-Strand Synthesis System* for RT-PCR (Invitrogen) and collected cDNA amplified using PCR. Identification of *_Ak_*VAL opsin gene was conducted via multi sequence alignment BLAST algorhythm with *A. katoptron* retinal transcriptome databank and matching results controlled with pairwise EMBOSS-Needle alignments with other *A. katoptron* opsin genes (for details of mRNA isolation, sequencing and identification of opsin transcripts see (Mark et al. 2018). Sequence alignments and creation of the phylogenetic tree were performed using Uniprot.

### Generation of plasmid constructs & mutagenesis

Generated *_Ak_*VAL opsin gene was cloned into the pAAV-CW3SL-eGFP vector (GenBank accession number: KJ411916.2) under the control of CMV promotor using *InFusion Aqua cloning (Beyer et al. 2015) kit* (Takara) to enable packaging capacities for viral production. For imaging purposes, we created an additional vector with the exchange of eGFP to the red fluorescent protein mCherry. All cAMP-based assays were performed using the *GloSensor* plasmid provided by the *GloSensor Technology* kit (Promega). The GsX Assay was executed with the additional use of Addgene provided chimeras of G_s_ (#109350), G_s_i (#109373), G_s_q (#109356), G_s_z (#109355), G_s_o (#109375), G_s_t (#109376), G_s_12 (#109357), G_s_13 (#109359) and G_s_15 (#109360). Control plasmid pAAV-CMV-mOpn4L was used unmodified as in previous publications (Spoida et al. 2016). Control plasmid pcDNA3.1 JellyOp was purchased from Addgene (#41432).

### Electrophysiological Patch Clamp Recordings

Human embryonic kidney (HEK) GIRK1/2 cells expressing GIRK1/2 subunits (provided by Dr. A. Tinker, UCL London) were constantly kept in the dark incubator at 37 °C and 5 % CO_2_ supplied with DMEM (Gibco) supplemented with 10 % FBS and G418 (5mg/ml). Cells were seeded into 35 mm plastic dishes 48 h prior and transfected with the respective plasmid using FuGene HD (Promega) lipofection 24 h prior to experiments. Under dark room conditions, immediately before Patch experiments the medium was exchanged to a high potassium external solution (20 mM NaCl, 120 mM KCl, 2 mM CaCl_2_, 1 mM MgCl_2_, 10 mM HEPES-KOH, pH 7.3 (KOH)) containing 1 µM *9-cis* (Sigma Aldrich) (if not explicitly stated otherwise). Patch pipettes with a resistance of 2-5 MΩ were filled with a physiological internal solution (40 mM KCl, 5 mM MgATP, 10 mM HEPES-KOH, 5 mM NaCl, 2 mM EGTA, 2 mM MgCl_2_, 0.01 mM GTP, pH 7.3 (KOH)). Experiments were performed with an inverted microscope (Leica), micromanipulators (Luigs & Neumann) and recordings displayed via the software PATCHMASTER. Manipulation of light pulse wavelength, duration and intensity were controlled by the monochromator system Polychrom 5 (TILL Photonics) and the Software PolyCon. In stated experiments dishes were incubated with 200 ng/ml PTX 24 hours prior to experiments to block G_i_ specific currents or 10 µM U73122 to inhibit PLC-dependent effects. More detailed experimental conditions can be looked up in previous publications (Spoida et al. 2016; Karapinar et al. 2021). Whole-cell Patch Clamp recordings were analyzed using Igor Pro software and exemplary traces smoothed every 5000 data points using the median value. Final statistics and graphical editing were performed using SigmaPlot (FromSoftware) and CorelDraw.

### 2-Photon Calcium Imaging

HEK293 tsA 201 cells were kept under identical conditions. Cells were seeded 48 hours prior and co-transfected with the respective opsin (mOpn4L-mCherry or *_Ak_*VAL-mCherry) and GCaMP6m 24 hours prior to the experiment. Imaging was performed in regular DMEM high glucose medium under dark room light conditions. Calcium signals were measured using a 2P-Setup (Bruker) using the 965 nm laser for the sensor excitation and Polychrom V monochromator system as the light source for additional opsin excitation with varying spectral properties. Cell surface level was adjusted using red light to exclude prior opsin activation. Acquired fluorescent data was normalized to their respective starting values and graphical design performed with SigmaPlot and CorelDraw.

### *In vitro* cAMP GloSensor and GsX Assay

HEK293 tsA 201 cells were kept as described above. Cells were seeded into poly-L-lysin coated 96-well plates 48 hours and transfected with the GloSensor, the respective opsin (*_Ak_*VAL, JellyOp) and for GsX experiments required chimeras (see Fig 3a) in a ratio of 100:100:1. The medium was exchanged to L-15 medium containing 2 mM beetle luciferin (Promega) and 1 µm *9-cis* Retinal (Sigma Aldrich) 1 hour before the experiment begins. Luminescent data was acquired every 40 s using a plate reader (PerkinElmer 2030 Multilabel reader VICTOR X3) and a counting time of 1 s. After baseline levels were recorded for 160 s, well plates were stimulated with 0.8 mW/cm^2^, 470 nm stimulation using a custom build LED device before post stimulation luminescent values were recorded for 1300 s. To artificially increase cAMP levels to examine adenylate cyclase activity reduction by G_i_ activation, wells were incubated in 3 µM Forskolin 30 minutes prior to the experiment. For GsX data, mean luminescent values were normalized to the mean starting value, corrected to negative control luminescent values and plotted in a radar plot indicating a 0,5-fold increase for each segment. For 3b luminescent data starting values were scaled to 1 showing the relative decline of luminescence.

### TRUPATH Assay (Olsen et al. 2020)

ΔG7 HEK293A (Bowin et al. 2019) were kept as described above and cultured in DMEM supplemented with 10% FBS and 1% Penicillin/Streptomycin (P/S) (Sigma Aldrich). Cells were seeded into clear bottom, white 96-well plates (Greiner Bio One), after being coated with poly-L-lysine (Merck Sigma Aldrich). The next day the medium was changed to DMEM/10%FCS/1%P/S containing 10 µM *9-cis*-Retinal (Sigma Aldrich). After this, plates were kept wrapped aluminum foil until the measurement. Cells were co-transfected with RLuc8-Gα, Gβ, Gγ (Addgene) and *_Ak_*VAL at a 1:1:1:1 ratio, a total of 100 ng DNA/well was transfected using Lipofectamine 2000 (Thermo Fischer). Cells were incubated for 24 h and the medium was then replaced with Hanks Balanced Salt Solution (HBSS) without phenol red (Capricorn) with 20 mM HEPES (ROTH), 10 µM *9-cis*-Retinal, and 50 µM Coelenterazine 400a (Nanolight Technologies) for 5 minutes at RT prior to the measurement. A BMG Clariostar Plus plate reader was used and cells were activated with 385 nm for 1 s using a coolLED pE 4000. Eight individual end point measurements were done immediately after one another to determine the activation of the receptor based on the ratio changes. The ratio changes were measured from RLuc8-Gα and Gγ-GFP2 emission with the filters 410-80 and 515-30, directly after light activation.

### *In vitro* cerebellar slice electrophysiological recordings

All experiments were performed under red light conditions. As described previously (Surdin et al. 2023) acute sagittal, cerebellar slices (250 µm) were prepared 6-8 days after AAV injection (Vibratome VT1000S, Leica) in a dissection solution ((in mM): 87 NaCl, 2.5 KCl, 0.5 CaCl_2_, 7 MgCl_2_, 1.25 NaH_2_PO_4_, 25 NaHCO_3_, 10 D-Glucose and 75 Sucrose enriched with 95% O_2_ and 5% CO_2_). Slices were incubated in recording solution ((in mM): 125 NaCl, 2.5 KCl, 2 CaCl_2_, 1 MgSO_4_, 1.25 NaH_2_PO_4_, 26 NaHCO_3_, and 20 D-Glucose enriched with 95% O_2_ and 5% CO_2_) at 37°C for 1h and stored at room temperature. Slices were preincubated for 10 min in recording solution containing 25 µM *9-cis*-retinal or ATRA, 0.025% (±)-α-tocopherol (Sigma), and 0.2% essentially fatty acid free albumin from bovine serum (BSA, Sigma-Aldrich) before recording. Glass pipettes (3 – 5 MΩ, MPC325, Sutter Instrument) were filled with internal solution ((in mM): 125 Potassium Gluconate, 10 HEPES, 4 NaCl, 2 MgCl_2_, 0.2 EGTA, 4 Mg-ATP, 0.4 Na-GTP, and 10 Tris-Phosphocreatine (pH 7.3, 280mOsm)). *_Ak_*VAL-eGFP expressing Purkinje cells were visually identified using a 40x objective attached to an upright microscope (BX51WI, Olympus) using 470 nm light stimulation (Polychrome V, TILL Photonics). Whole-cell recordings were low pass-filtered at 3 kHz using an EPSC10 amplifier (HEKA) and conducted on cells that were voltage clamped to −60 mV. To measure light-induced and opsin-mediated effects on neuronal activity cells were held at a constant base current that elicited moderate spontaneous activity before a train of 10 s different light pulse conditions was applied (light off/ 560nm/light off/380 nm/light off). To measure membrane potential, rheobase, spike rate, latency to first induced AP and input resistance the response to different currents steps was recorded (−200 to 300 pA, step 100 pA, 1s current step, 2s repetition rate). PatchMaster software (HEKA) was used for data acquisition and MATLAB for ofline analysis.

### Protein expression and purification

A cDNA coding for the full length *_Ak_*VAL sequence was codon optimized for insect cells expression and cloned into a pFASTBAC1 vector following an N-termini human influenza hemagglutinin (HA) membrane localization tag and a FLAG peptide. Baculoviruses, prepared according to the Bac-to-Bac™ manufacturer protocol (Invitrogen), were used to infect *Spodoptera frugiperda* (Sf9) insect cells at 100x viral dilutions. Cells were cultured in ESF921 medium and infected at a density of 4 million cells/mL. Expression was carried out for 60 h at 27 °C and cells were harvested by centrifugation and stored at −80 °C. Cells were lysed at 4 °C for 1 h using a lysis buffer containing 20 mM TRIS-HCL (pH 7.5), 1 mM Ethylenediaminetetraacetic acid (EDTA), 100 μM tris(2-carboxyethyl)phosphine (TCEP), 100 μM *9-cis* retinal (Merck), 15% (W/V) glycerol and protease inhibitor cocktail (PIC) containing 1 mM phenylmethylsulfonyl fluoride (PMSF), 160 µg/mL benzamidine and 2.5 µg/mL leupeptin. Membrane harvesting was done under centrifugation at 37,000 x g for 30 min at 4 °C. Cell pellets were homogenized in a Dounce homogenizer with solubilization buffer containing 20 mM HEPES-NaOH (pH 7.5), 150 mM NaCl, 1 % (w/v) n-Dodecyl-β-D-maltoside (DDM), 0.1 % (w/v) cholesteryl hemisuccinate (CHS), 100 μM TCEP, 15% (W/V) glycerol and PIC. Solubilization was carried out under gentle agitation for 1.5 h at 4 °C. Non-solubilized material was removed by centrifugation at 37,000 x g for 30 min at 4 °C. Clarified supernatants were loaded onto an anti-FLAG column. The FLAG-beads (GeneScript) were pre-equilibrated with 3 column volumes (CVs) of wash buffer containing 20 mM HEPES-NaOH (pH 7.5), 150 mM NaCl, 0.1 % (w/v) DDM, 0.01 % (w/v) CHS and 100 μM TCEP. Following sample loading, the beads were washed with 20 CVs of wash buffer, and eventually the protein was eluted with an elution buffer containing the same ingredients as the wash buffer supplemented with 0.25 mg/mL FLAG peptide (DYKDDDDK peptide, GeneScript). The protein was collected and concentrated in a 100 kDa molecular weight cut-off (MWCO) concentrator to ∼8 mg/mL and was kept frozen in the dark in −80 °C until further use. Purification has been done in a dark room, with an occasionally dim red light.

### Biophysical absorption measurements

Optical measurements were carried out using the wavelength scan program in a nanophotometer (Implen™ NanoPhotometer™ NP80). Following a first λ scan in the UV-vis spectra for a *9-cis* reconstituted sample prepared in the dark, two types of experiments were conducted: (1) tandem illumination at wavelength that corresponds to the maximal absorbance of the sample, by alternate illuminations at green (520 nm) and UV (390-400 nm)/blue (450 nm) lights. The experiment was done at 4 °C, illumination was allowed for 30 sec for the UV and 1 min for the blue/green lamps. λ scan was taken right after each respective illumination. An overall of 5 alternate illuminations were carried out repetitively. And (2) thermostability measurement, by which the sample was kept at room temperature (25 °C) for 10 minutes in the dark after each illumination treatment. λ scan was taken after each cycle of illumination and dark incubation period. The assay was repeated 5 times. Difference spectra were calculated by substituting the OD values of sequential samples illuminated with a certain wavelength (e.g., green) from the previous light treatment (e.g., dark or UV illumination). Values are normalized to baseline recorded beyond 700 nm.

### In vivo *C. elegans* behavioral experiments

Caenorhabditis elegans strains were kept on nematode growth medium (NGM) seeded with Escherichia coli strain OP50-1 as described previously (10.1093/genetics/77.1.71). The strains KG1180 (lite-1(ce314)) and MT363 (goa-1(n363)) were provided by the Caenorhabditis Genetics Center (CGC, University of Minnesota, USA). Transgenic animals were generated by microinjection of pCR09 (punc-17:: *_Ak_*VAL opsin::mCherry) into the gonad of young adult animals as described previously (Fire 1986).

Confocal images of *_Ak_*VAL opsin::mCherry expression were acquired with a LSM 780 confocal microscope system equipped (Zeiss, Germany) with a Plan-Apochromat 63 x oil objective using the Zeiss ZEN software tile scanning module. Young adult animals were imaged on 7 % agarose pads and anesthetized with 20 mM tetramisole hydrochloride (Sigma Aldrich, USA). Fluorescence images were processed with ImageJ (Schindelin et al. 2012).

All experiments were conducted in a *lite-1* knockout background to avoid triggering the endogenous reaction of *C. elegans* to light (Edwards et al. 2008). Unstarved, transgenic L4 larvae were picked ∼ 18 h before the experiment. *9-cis* retinal (Sigma Aldrich, USA) was supplied to the animals by adding 100 µM to the OP50 suspension before seeding of NGM plates. Experiments were performed on three different days with animals picked from different populations and animals were kept in darkness until assessment. Measurements using different conditions were done on the same day. Crawling speed was measured as described previously (Vettkötter et al. 2022; Seidenthal et al. 2023). Briefly, animals were washed three times with M9 buffer (K2PO4 20 mM; Na2HPO4 40 mM; NaCl 80 mM; MgSO4 1 mM) to remove residual OP50 before being transferred to unseeded NGM plates. Animals were allowed to settle for 15 minutes and videos were acquired using a Falcon 4M30 camera (DALSA, USA). Custom-build LED rings (Alustar 3W 30°, ledxon, 470 nm for blue light, MinoStar grün Optik 30°, Signal-Construct, 525 nm for green light and UV SMD-LED 3535 1.8W 55°, LG Innotek, 365 nm for UV light stimulation, respectively) controlled by an Arduino Uno (Arduino, Italy) device operating on a custom-written Arduino script were used to apply light stimulation. Crawling data was analyzed using the multi-worm-tracker “Choreography” software (Swierczek et al. 2011) and summarized using a custom-written Python script. Crawling speed was normalized to the average before stimulation. Measurement of body lengths was performed as described previously (Seidenthal et al. 2022). In short, videos of manually tracked, individual animals were acquired on unseeded NGM platesStimulation light from a 50 W HBO was applied through a 380/14 Brightline HC excitation filter (AHF Analysentechnik, Germany). Body length was calculated using the WormRuler software by normalization of skeleton lengths to the average before stimulation (Seidenthal et al. 2022). Values which were higher than 120 % or lower than 80 % of the initial body length were discarded as they are likely to represent artifacts resulting from the background correction (e. g. coiling of animals).

